# The archaeal class *Nitrososphaeria* is a key player of the reproductive microbiome in sponges during gametogenesis

**DOI:** 10.1101/2024.07.03.601946

**Authors:** Marta Turon, Vasiliki Koutsouveli, María Conejero, Sergi Taboada, Aida Verdes, José María Lorente-Sorolla, Cristina Díez-Vives, Ana Riesgo

## Abstract

Sponge-associated microbes play fundamental roles in regulating their hosts’ physiology, yet their contribution to sexual reproduction has been largely overlooked. Most studies have concentrated on the proportion of the microbiome transmitted from parents to offspring, providing little evidence of the putative microbial functions during gametogenesis in sponges. Here, we use 16S rRNA gene analysis to assess whether the microbial composition of five gonochoristic sponge species differs between reproductive and non-reproductive individuals and correlate these changes with their gametogenic stages. In sponges with mature oocytes, reproductive status did not influence either beta or alpha microbial diversity. Interestingly, in two of the studied species, *Geodia macandrewii* and *Petrosia ficiformis,* which presented oocytes at the previtellogenic stage, significant microbial composition changes were detected between reproductive and non-reproductive individuals. These disparities were primarily driven by differentially abundant taxa affiliated with the *Nitrososphaeria* archaeal class in both species. We speculate that the previtellogenic stages are more energetically demanding, leading to microbial changes due to the phagocytosis of microbes to meet nutritional demands during this period. Supporting our hypothesis, we observed significant transcriptomic differences in *G. macandrewii*, mainly associated with the immune system, indicating potential changes in the sponge’s recognition system. Overall, we provide the first insights into the possible roles of sponge microbiomes during reproductive periods, potentially uncovering critical interactions that support reproductive success.

## INTRODUCTION

All animals on Earth engage in interactions with environmental microbes, leading to the evolution of complex symbioses that range from benign to pathogenic (M. J. McFall-Ngai 2002). These microorganisms have coevolved with their hosts in such a way that holobionts – multicellular eukaryotes together with their populations of persistent symbionts (Margulis, L. 1991) – are considered a unit of selection, as postulated by the hologenome evolutionary theory (Zilber-Rosenberg and Rosenberg 2008). Since microbes can significantly influence host biology and speciation (Shropshire and Bordenstein 2016; M. McFall-Ngai et al. 2013), natural selection may act at the holobiont level rather than at the species level (Dittami et al. 2021). While there is mounting evidence that microbiomes affect host health, physiology, development and behaviour (Rowe et al. 2020), their effects on host reproduction have only recently begun to be considered (Comizzoli et al. 2021).

The reproductive microbiome comprises bacteria, viruses, unicellular algae, protozoans, and fungi living in or on any structure, organ, fluid, or tissue of a host that typically comes into contact with the gametes, reproductive tract or organs of another individual through mating, spawning, or pollination (Rowe et al. 2020; Comizzoli et al. 2021). These microbiomes have the potential to significantly impact host fertility and fitness, with implications for sexual selection, sexual conflict, mating systems, and reproductive isolation (Rowe et al. 2020). While reproductive microbiomes may have a direct role in reproduction, microbial communities from other body parts can also play an indirect role in regulating reproductive processes. Although these effects are well recognized in humans and model organisms, they remain poorly studied in wild animal species, especially invertebrates (Comizzoli et al. 2021).

To date, the most striking example of the effect of microbial symbionts on the reproductive biology of their hosts is provided by the *Wolbachia* endosymbiont and its variety of arthropod hosts. These symbionts manipulate host reproduction to enhance their inheritance through the female germline, primarily through cytoplasmic incompatibility (Beckmann, Ronau, and Hochstrasser 2017; LePage et al. 2017; Shropshire et al. 2018). This means that embryos are only viable if both partners are infected. Furthermore, the presence of *Wolbachia* has been shown to induce parthenogenesis in mites (Weeks and Breeuwer 2001). In aquatic systems, certain microbial symbionts have been demonstrated to enhance reproduction in the pennate diatom *Seminavis robusta* (Cirri et al. 2019), the marine diatom *Odontella* sp. (Sison-Mangus et al. 2022), the moon jellyfish *Aurelia aurita* (Weiland-Bräuer et al. 2020) and in the freshwater polyp *Hydra viridis* (Habetha et al. 2003). In all of these examples, bacteria played a critical role in promoting the initiation of reproduction. Their absence resulted in reduced reproductive success, either by decreasing the production of bacteria-derived sexual pheromones or by failing to induce oogenesis. Beyond the manipulation of host reproduction triggering their gametogenesis, very little is known about the dynamics of the microbiota during the essential period of sexual reproduction in aquatic invertebrates. For instance, the slight changes in microbiota composition reported between male and female corals (Wessels et al. 2017), and the large changes in the abundance of Rickettsiales between sexual and asexual types of freshwater snails (Takacs-Vesbach et al. 2016), both point to a link between reproductive mode and the bacterial microbiome in invertebrates that has been scarcely studied.

Sponges are among the aquatic animals that exhibit some of the most prominent and diverse symbiotic partnerships (Webster and Thomas 2016). Their microbial counterparts play vital roles in nutrient acquisition, chemical defence and host physiology, underscoring their importance in maintaining sponge health and ecological success (Taylor et al. 2007). However, while the functional roles of these microbes during reproduction have been hypothesized (Díez-Vives et al. 2022), they have rarely been demonstrated. Sponges display a wide range of reproductive strategies, including asexual and sexual reproduction, hermaphroditism, gonochorism, sequential hermaphroditism, oviparity, and viviparity. Research on sponge reproduction has primarily focused on how microorganisms are transmitted and incorporated into the germline, as microbial composition and structure typically changes during the life cycle of sponges (Carrier et al. 2022; Díez-Vives et al. 2022; Engelberts et al. 2022). However, there is limited evidence on the specific roles of these microbes during these developmental processes. For instance, the interplay between the larval molecular toolkit and the metabolites produced by their symbiotic bacteria is essential for inducing settlement in a marine sponge (Song, Hewitt, and Degnan 2021). But what happens during gametogenesis has not been actively researched. Díez-Vives et al. (2022) suggested that changes in the sponge-microbial cross–talk are likely to occur during gametogenesis in gonochoristic sponges. During spermatogenesis, part of the choanocyte chambers are transformed into spermatic cysts (Maldonado and Riesgo 2009), compromising the sponge’s filtering capacity and affecting its nutrition. On the other hand, oogenesis requires a high nutritional demand due to the significant amount of yolk produced for the eggs. Therefore, microbes might play crucial roles in meeting the nutritional demands required during these energetic processes to ensure the survival of individuals. Evidence of symbiont digestion during reproduction has been reported (Usher et al. 2001; Oren, Steindler, and Ilan 2005), but whether digestion of symbionts occurs randomly or selectively remains to be elucidated. In that latter scenario, a selective digestion of symbionts would imply changes in the final microbial composition of the sponge during reproduction.

In the present study, we investigate the microbial composition of five gonochoristic sponge species during their sexual reproductive period to determine whether there are changes in the microbiome composition and structure between reproductive and non-reproductive individuals of the same species and/or between females and males. Using 16S rRNA gene analysis, we evaluate whether these species exhibit differentially abundant microbes during reproduction and correlate these microbial changes with the gametogenic stages observed through histological analysis. Additionally, for a single species, we study the host transcriptional responses during reproduction with the aim to detect enriched functions that might be related to the putative microbiome changes. Overall, we aim to provide the first insights into the possible roles of sponge microbiomes during reproduction to elucidate critical interactions that may be supporting reproductive success.

## METHODS

### Sample Collection, DNA extraction and 16S rRNA gene amplification

We collected 106 sponge specimens from five species, during various sampling campaigns (Table S1). These species encompassed four High Microbial Abundance (HMA) sponges, namely *Geodia macandrewii* (n = 6)*, Geodia hentscheli* (n = 14)*, Petrosia ficiformis* (n = 23) and *Chondrosia reniformis* (n = 58), as well as one Low Microbial Abundance (LMA) sponge, namely *Topsentia* sp. (n = 5). Samples of *G. macandrewii* were collected in the Norwegian fjords (Korsfjord, 59°58.8790’ N, 5°22.4371’ E) at 80 m during September 2016, and those of *G. hentscheli* were collected in two different sites in the Vesterisbanken seamount (at mesobenthic depths=140 m: 73°31’10.0” N 9°09’36.0” E and at deeper sites (500-700 m): 72°40’51.964” N, 2°51’44.481” E) in August, 2019. The individuals of *P. ficiformis* were collected in two different Mediterranean sites: Naples, Italy (40°49’16.5” N 14°04’39.8” E) in July, 2022 and in L’Escala, Spain (42°7’36.369” N 3°7’56.337” E) in December, 2022. Samples of *Chondrosia reniformis* were collected in Santa Anna beach, Blanes, Spain (42°7’36.369” N 3°7’56.337” E) in July of two different years, 2020 and 2021. Finally, *Topsentia* sp. was collected in the Cantabrian Sea (43°52.312’ N, 5°54.106’ W) at 695 m, in June 2017.

Each individual sponge specimen was subjected to dual preservation methods: tissue fragments of ca. 1 cm^3^ were preserved in 4% formaldehyde in seawater and/or 2.5% glutaraldehyde in PBS or 0.4M PBS and 0.34M NaCl for subsequent histological observations, while additional tissue fragments of ca. 1 cm^3^ were preserved in either absolute ethanol or RNAlater and frozen at –20 °C until subsequent nucleic acid extractions.

DNA was isolated per sample using a Dneasy Blood and Tissue kit (Qiagen, manufacturer’s protocol, July 2006 version). The 16S rRNA gene V4 region was amplified using the universal microbial primers 515F-Y (Parada, Needham, and Fuhrman 2016) and 806R (Apprill et al. 2015), which amplify both bacterial and archaeal members. DNA amplification was always performed in duplicate, and libraries were prepared with the Nextera XT DNA Library Preparation Kit (Illumina Inc.). Next-generation, paired-end sequencing was performed on the Illumina MiSeq platform at the ‘Genomics Unit’ of the Universidad Complutense de Madrid using v3 chemistry (2 x 300 bp).

### Assessment of reproductive state

Samples in formalin or were used to assess the reproductive status of the sponge individuals (sex (female/male), and gametogenic stage. All samples with siliceous spicules (i.e. *Geodia macandrewii, Geodia hentscheli, Petrosia ficiformis*, and *Topsentia* sp.) were then desilicified in 5% hydrofluoric acid overnight and then rinsed with distilled water at least twice. The tissue was then processed for either light microscopy or Transmission Electron Microscopy (TEM). For light microscopy, tissues were dehydrated through an increasing ethanol series and later embedded in paraffin after a brief rinse in xylene. For TEM, sponge tissues preserved in glutaraldehyde were rinsed in PBS and 0.6 M NaCl, then post fixed in 2% osmium tetroxide in 0.4M PBS and potassium–ferrocyanide for 1–2 hrs at 4°C, and thoroughly rinsed in distilled water afterwards. As with samples for light microscopy, an increasing ethanol series was used to dehydrate the tissue for further embedding in epoxy resin.

The samples prepared for light microscopy were sectioned at 5 μm with an HM 325 rotary microtome (ThermoFisher-Scientific) and stained them with hematoxylin and eosin, using standard protocols, and mounted in slides with DPX. Slides were observed with an Olympus microscope (BX43) with a UC50 camera at the Museo Nacional de Ciencias Naturales de Madrid (CSIC). To assess the percentage of maturation of oocytes, the size (maximum diameter) of 30–50 oocytes for each sample were measured, and then compared to the known diameter of mature oocytes of these species (Riesgo and Maldonado 2008; Maldonado and Riesgo 2009; Koutsouveli et al. 2020).

For TEM, we obtained ultrathin sections (65 nm) from epoxy blocks with an Ultracut ultramicrotome (Reichert-Jung) and then used a uranyl acetate/lead citrate staining protocol (Reynolds, 1963). Ultrathin sections on gold coated grids were observed with a Hitachi Transmission Electron Microscope (H-7650) at the imaging facilities in Kew Botanical Gardens in the UK or at the JEOL JEM 1010 at the Centro Nacional de Microscopía Electrónica in Spain. The purpose of TEM was to observe phagocytosis of microbes by sponge cells in both reproductive and non-reproductive individuals.

### Microbiome pipeline analysis

Raw 16S rRNA gene sequences were processed separately using the UPARSE pipeline (Robert C. Edgar 2016). Briefly, primer sequences were removed and overlapping paired reads were merged using PEAR (J. Zhang et al. 2014). After quality check and de-replication, unique read sequences were detected using the *fastx_uniques* command. Denoising (error-correction) of amplicons was performed using *unoise3* (Robert C. Edgar 2016), which removed chimeras, reads with sequencing errors, PhiX, and low complexity sequences due to Illumina artefacts, and generated ZOTUs (“zero-radius” Operational Taxononic Units) with 100% identity sequences. Taxonomic assignment was done with SINA v1.2.11 (Quast et al. 2012) using SILVA 138 database. Sequences with low alignment quality (<75%) or unclassified as well as sequences identified as mitochondria or chloroplasts were removed from the analysis. The initial ZOTU table underwent rarefaction, with the minimum threshold being set at the sample containing the fewest reads, which amounted to 12,000 reads. This rarefaction process was performed using the *rrarefy* function from the R package *vegan* v2.5-6 (Oksanen, J. et al. 2019). The rarefied ZOTU table was subsequently employed for conducting alpha diversity analyses. Moreover, a relative abundance ZOTU table was utilized for characterizing the microbial community and conducting beta diversity analyses.

### Microbiome statistical analyses

We conducted a distance-based multivariate analysis of the microbial communities within the sponge samples at the ZOTU level using the *vegan* v2.5-6 (Oksanen, J. et al. 2019) in R (R Core Team 2020). To assess the dissimilarity between samples, we computed a Bray-Curtis dissimilarity distance matrix based on the log2-transformed relative abundance ZOTU table, which was utilized to visualize the patterns of microbial community composition in different sponge species. This visualization was achieved through the construction of a cluster dendrogram using the *hclust* function and a Principal Coordinates Analysis (PCoA) using the *cmdscale* function of the *vegan* R package. We first tested the effect of host identity (species) on the composition of microbial communities with non-parametric Permutational Analysis of Variance (PERMANOVA) using the *adonis* function using 999 permutations and a significance cut-off for p-values of 0.05. Following the evaluation of host identity’s impact on microbial communities, we created distinct subsets for each sponge species and collection site to examine the influence of the reproductive state on each subset. For each species, we calculated Bray-Curtis dissimilarity distance matrix and visualized the ordinations through PCoAs, as described above. The effect of reproduction (reproductive *vs* non-reproductive) and reproductive state (oocytes, sperm or non-reproductive) on the sponge microbiomes was tested using the *adonis* function. Distance to centroid (Betadispersion) between reproductive and non-reproductive individuals was calculated for each species, and analyses of variance (ANOVA) were conducted to test for significant differences. Moreover, generalized linear models were performed in the R package “EdgeR” v.3.26.8 (Robinson, McCarthy, and Smyth 2010) to discern variations in the abundance of particular microbiome ZOTUs across different reproductive stages. Heatmaps and bubble plots were used to visualize the differentially abundant ZOTUs between reproductive and non-reproductive individuals using *heatmaply* (Galili et al. 2018) and *ggplot* (Wickham 2016) R packages. A rarefied dataset was used to compute alpha diversity indices, specifically Richness, Shannon and inverse Simpson indexes, within the *vegan* package. Normality was checked using *Shapiro* test prior to perform an ANOVA or Kruksal-Wallis test to compare alpha diversity measures between reproductive and non-reproductive individuals for each species.

The evolutionary relationships of a subset of the most differentially abundant ZOTUs found in *G. macandrewii* and *P. ficiformis* (from Naples) belonging to the phyla *Crenarchaeota* and *Nitrosphaeria* class were investigated by constructing a phylogenetic tree with the most similar archaeal taxa retrieved by BLAST (Altschul 1997). An alignment of 79 archaeal sequences retrieved from NCBI and 47 ZOTUs from *G. macandrewii* and *P. ficiformis* was done with MUSCLE (R. C. Edgar 2004), containing 253 bp of the 16S ribosomal gene. The phylogenetic relationships were explored with RAxML-NG 1.1.0 (Kozlov et al. 2019) using the evolutionary model (JC) defined by ModelTest-NG v0.1.7 (Darriba et al. 2020), with transfer bootstrap expectation parameters, 10 runs and 100 replicates.

### Host Differential Gene Expression Analyses

To understand changes in the immune system of reproductive species, we used the reference transcriptome and raw reads from the published study by Koutsouveli et al. (2020) for *G. macandrewii*. We assessed the mapped read abundance using RSEM 7/2/2024 7:55:00 AM and then looked for differential gene expression (DE) between reproductive (pooling together males and females) and non-reproductive specimens with edgeR (Robinson, McCarthy, and Smyth 2010). We then used blast and BLAST2GOPRO to annotate our differentially expressed genes and used Gprofiler (Reimand, Arak, and Vilo 2011) against the human database to perform Gene Ontology (GO) enrichments (with Benjamini-Hochberg FDR corrections of 0.05). Those GO terms that passed the threshold were visualized with REVIGO using the associated corrected p-value.

## RESULTS

### Assessment of reproductive stages

Between 40-67% of the individuals in each population were engaged in reproduction (Table 1), with 34 female individuals showing oocytes in different stages of vitellogenesis (from mid-stage to fully mature), and 18 individuals showing sperm (Table 1, Figure 1). In most species engaged in oogenesis, females already had completed the process of yolk formation and showed mature, vitellogenic oocytes (Table 1), while only two species were actively involved in yolk formation: *G. macandrewii* and *P. ficiformis* from Naples (Figure 1 & Figure 2, Table1). Our TEM micrographs showed phagocytosis of microbial cells found in the mesohyl (symbionts), mostly in reproductive specimens of *G. macandrewii*, *P. ficiformis*, and *C. reniformis* (Figure 2). In particular, both archaeocytes and oocytes of *G. macandrewii* female specimens were observed phagocytosing microbial cells to make yolk platelets (Fig. 2A–B), although in the case of oocytes, many of the endocytosed microbes were not digested but remained stored in vesicles product of vertical transmission of the microbiota (Fig. 2B). In *P. ficiformis*, the nurse cells that were aggregated around the oocyte showed large vesicles with evidences of microbial digestion and yolk platelets, which were later transferred to the oocytes (Fig. 2C). Similarly, archeocytes were observed close to the bacteriocytes with large vesicles containing digested bacteria (Fig. 2D). In *C. reniformis*, both nurse cells and archaeocytes were also observed phagocytosing microbes from the mesohyl for yolk formation (Fig. 2E–F).

**Figure 1.**
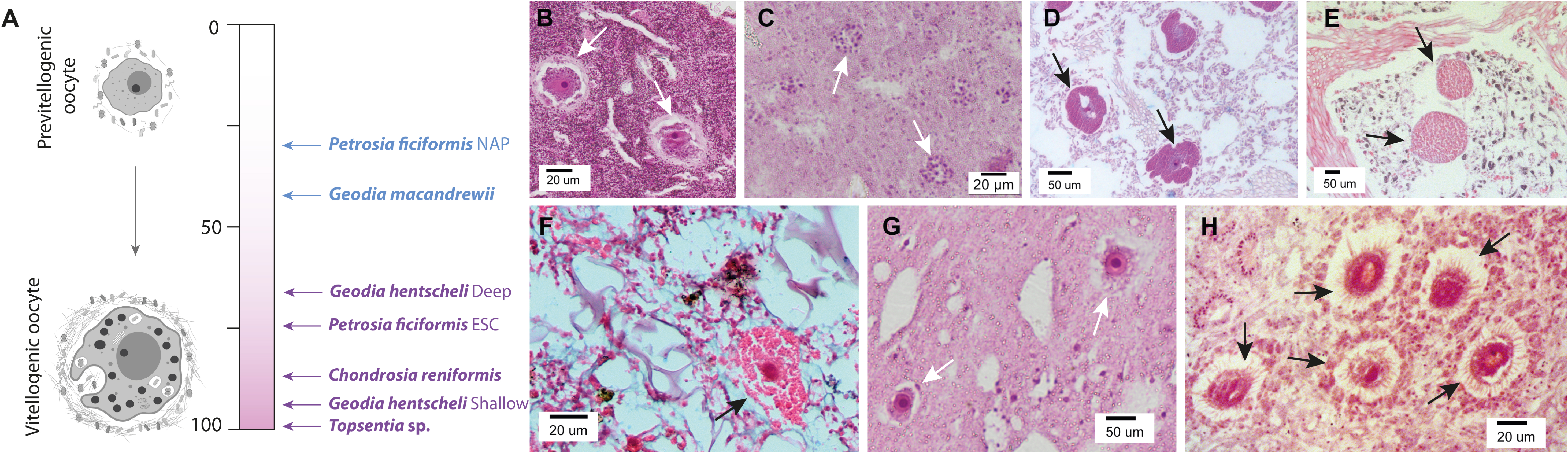
Oogenesis of the sponge species. **A**. Maturation gradient during oogenesis from previtellogenic to vitellogenic (mature) oocytes in the different species studied here. **B.** Previtellogenic oocytes of *Geodia macandrewii*. **C.** Developing spermatic cysts in *Geodia macandrewii*. **D.** Previtellogenic oocytes in *Petrosia ficiformis* from Naples (NAP). **E.** Vitellogenic oocytes in *Petrosia ficiformis* from L’Escala (ESC). **F.** Vitellogenic oocyte in *Topsentia* sp. **G.** Vitellogenic oocytes of *Geodia hentscheli*. **H.** Vitellogenic oocytes of *Chondrosia reniformis*. Arrows always indicate gametes.

**Figure 2.**
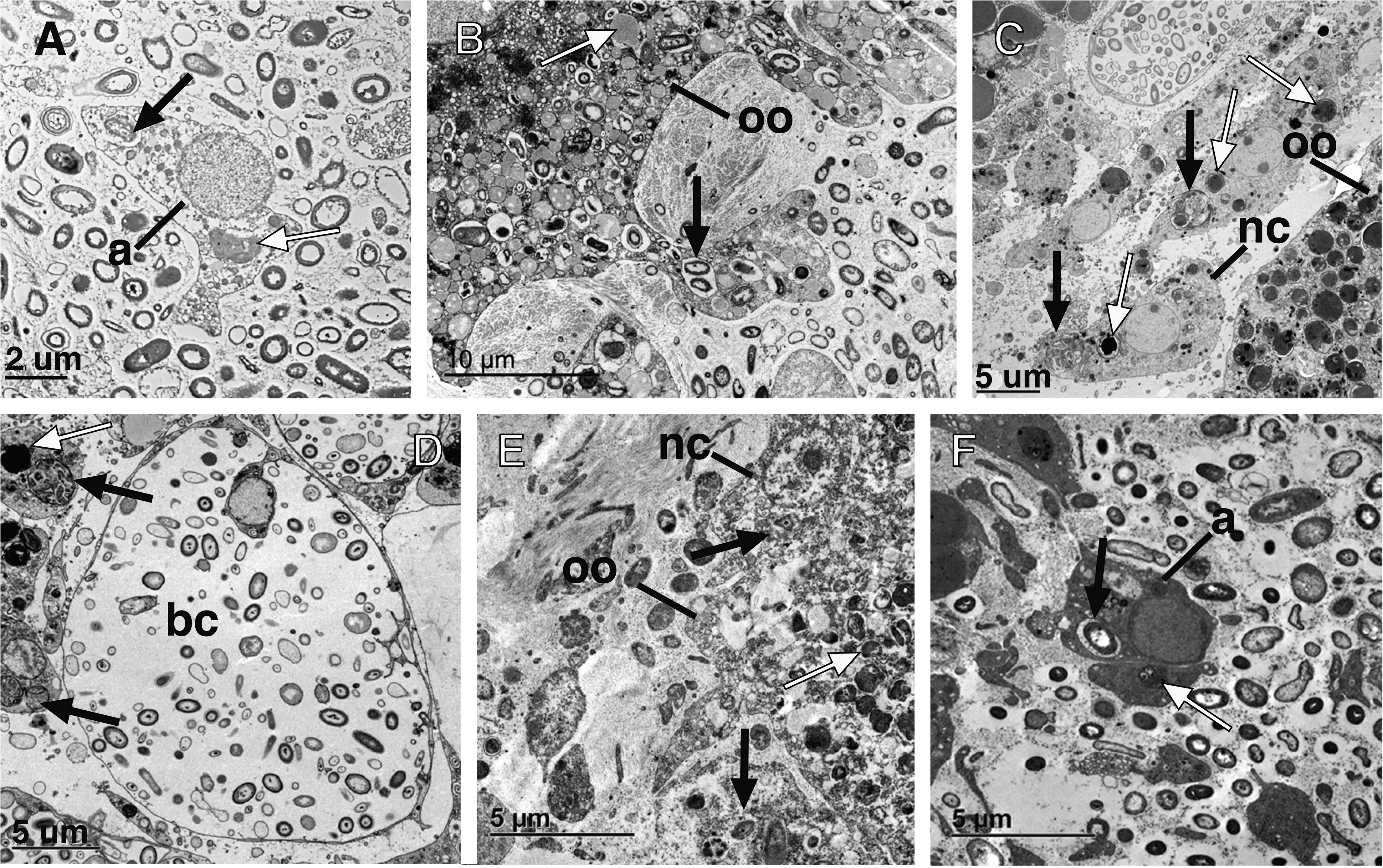
Ultrastructural images of microbial phagocytosis in sponges. **A.** Archaeocyte (a) phagocytosing microbes (black arrow) and making yolk (white arrow) in a female specimen of *Geodia macandrewii*. **B.** Detail of the phagocytotic processes of microbes (black arrow) occurring in previtellogenic oocyte projections (oo) of *Geodia macandrewii* and the yolk (white arrow) resulting from the phagocytosis. **C.** Periphery of the oocyte (oo) in *Petrosia ficiformis* during vitellogenesis, showing abundance of nurse cells (nc) with vesicles full of phagocytosed microbia (black arrow) and producing yolk (white arrow) that is later transferred to the oocyte (oo). **D.** Bacteriocytes (bc) and archaeocytes showing vesicles full of phagocytosed microbia (black arrow) and producing yolk (white arrow) in a male specimen of *Petrosia ficiformis*. **E.** Vitellogenic oocyte (oo) of *Chondrosia reniformis* with a nurse cell (nc) in the periphery phagocytosing microbial cells (black arrow) which are later transferred to the oocyte as yolk platelets (white arrow). **F**. Archaeocyte (a) phagocytosing microbes (black arrow) and making yolk (white arrow) in a female specimen of *Chondrosia reniformis*.

**Table 1.**
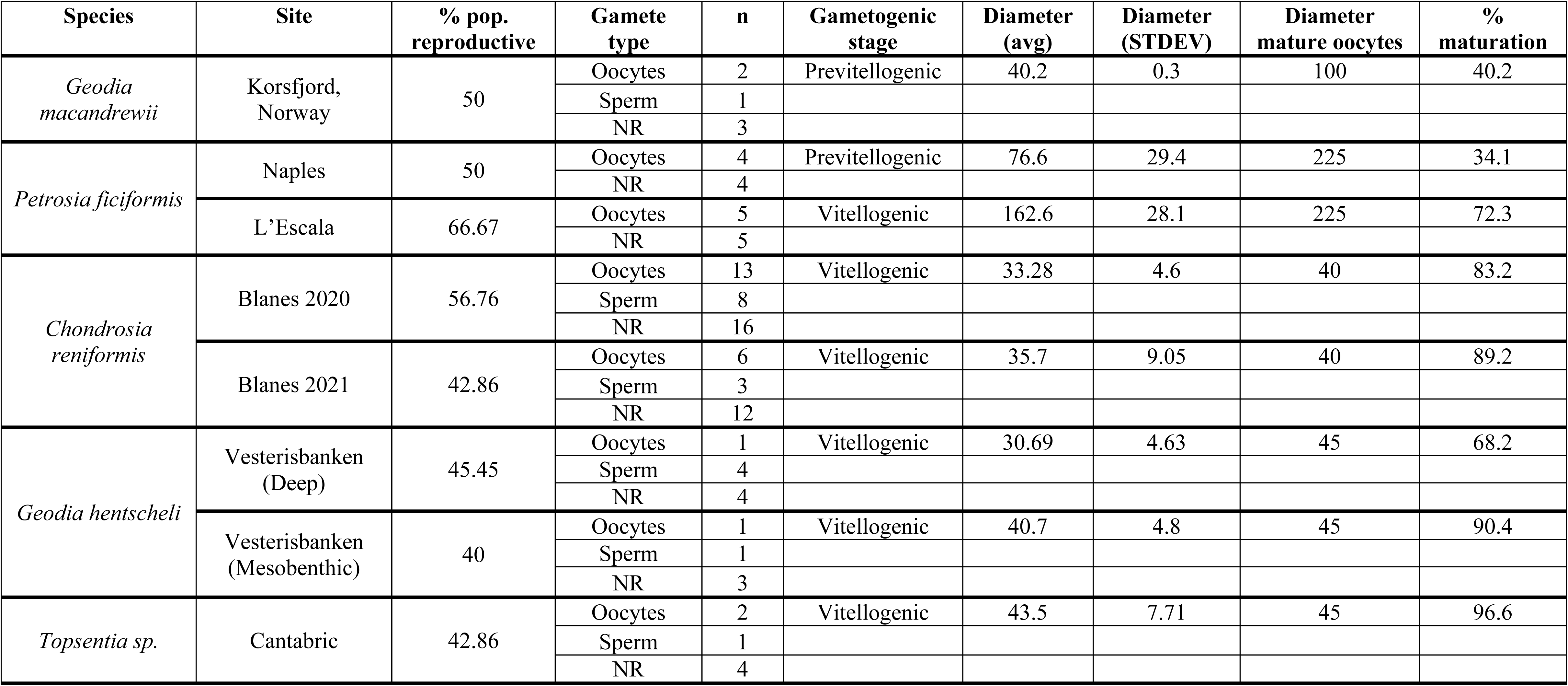
Information about reproductive stage of the sponge species used in this study. For each species and location, we show the percentage of the population involved in reproduction, the number of replicates in each stage, and the average size and maturation stage of the oocytes. NR = Non reproductive.

### Taxonomic composition and ordination of sponge microbial communities

Within the 106 samples belonging to 5 different sponge species sequenced for this study, we identified a total of 3,829 ZOTUs. The number of ZOTUs per species ranged from the lowest values of 536 ZOTUs in *C. reniformis* from Blanes, to the highest number of 2168 ZOTUs in *Topsentia* sp. (Table 2). These ZOTUs belonged to 37 different prokaryotic phyla (Table S2) with *Crenarcheota* being the most abundant phylum across samples, with an average relative abundance (avgRA) of 24.1%, followed by *Proteobacteria* (20.7%) and *Chloroflexi* (20.1%). At the class level, 75 different classes were identified (Table S3), with *Nitrososphaeria* being the most dominant class, with an avgRA of 24.1%, followed by *Gammaproteobacteria* (13.0%) and *<u>Alphaproteobacteria</u>* (11.1%).

**Table 2.** Differential Abundance (DA) analysis for each species and location. Column 5-7 display the results DA analysis when comparing reproductive versus non-reproductive individuals, while columns 8-9 present the results when comparing only oocytes versus non-reproductive individuals. The “n” values indicate the number of replicates.

| Species | Site | n | Num. ZOTUs | State | n | DA ZOTUs | State | n | DA ZOTUs |
| --- | --- | --- | --- | --- | --- | --- | --- | --- | --- |
| *Geodia macandrewii* | North Atlantic | 6 | 1545 | Repro | 3 | 156 | oocytes | 2 | 144 |
|  |  |  |  | NR | 3 | 239 | NR | 3 | 217 |
| *Petrosia ficiformis* | Naples | 8 | 835 | Repro | 4 | 13 | oocytes | 4 | X |
|  |  |  |  | NR | 4 | 15 | NR | 4 |  |
|  | L’Escala | 15 | 936 | Repro | 10 | 0 | oocytes | 5 | 0 |
|  |  |  |  | NR | 5 | 4 | NR | 5 | 0 |
| *Chondrosia reniformis* | Blanes | 37 | 536 | Repro | 21 | 0 | oocytes | 13 | 1 |
|  |  |  |  | NR | 16 | 2 | NR | 16 | 0 |
|  | Naples | 21 | 738 | Repro | 9 | 5 | oocytes | 6 | 1 |
|  |  |  |  | NR | 12 | 0 | NR | 12 | 0 |
| *Geodia hentscheli* | Vesterisbanken (Deep) | 9 | 1065 | Repro | 5 | 0 | oocytes | 1 | X |
|  |  |  |  | NR | 4 | 5 | NR | 4 |  |
|  | Vesterisbanken (Mesopelagic) | 5 | 957 | Repro | 2 | 1 | oocytes | 1 | X |
|  |  |  |  | NR | 3 | 6 | NR | 3 |  |
| *Topsentia sp.* | Cantabric | 5 | 2168 | Repro | 3 | 1 | oocytes | 2 | 0 |
|  |  |  |  | NR | 4 | 24 | NR | 4 | 14 |

Host identity was the main factor structuring the sponge microbiomes (ANOVA, *p* < 0.001), with a clear separation between HMA and LMA species (Figure 3A). Among the HMA species, *Nitrososphaeria, Dehalococcoidia, Gammaproteobacteria* and *Vicinamibacteria* emerged as the most prominent classes, displaying varying abundances across the four species (Figure 3B, Table S3). Conversely, the LMA species, *Topsentia* sp. was characterized by the prevalence of *Nitrososphaeria* (∼36% AvgRA) followed by *Gammaproteobacteria* (∼25% avgRA). In *Topsentia* sp. we also detected the presence of *Rickettsiales,* although at very low abundances (< 0.1 % avgRA), which did not display significant differences between reproductive and non-reproductive individuals. Taxonomic composition at class level remained consistent irrespective of sponge’s location or reproductive stage (Figure 3B). However, at the finer ZOTU level, the geographic location significantly separated the microbial communities of the sponge species with individuals in two different sites (Figure 3C, Supplementary Stats). Similarly, alpha diversity values varied significantly among the analysed sponge species (Table S1), with notable distinctions revealed by the Shannon and inverse Simpson indexes (ANOVA, p < 0.001, Supplementary Stats). Shannon diversity indexes, ranged from 5.39 in an individual of *Topsentia* sp. to 3.2 in an individual of *C. reniformis* (Table S1). Regarding the inverse Simpson index, the lowest value (7.7) was observed in an individual of *G. hentscheli*, while the highest value (84.23) was found in *P. ficiformis* (Table S1). The substantial disparities in alpha and beta diversity measures across the sponge species prompted the adoption of separate analyses for each sponge species and location to assess the influence of reproductive stage on the sponge microbiomes.

**Figure 3.**
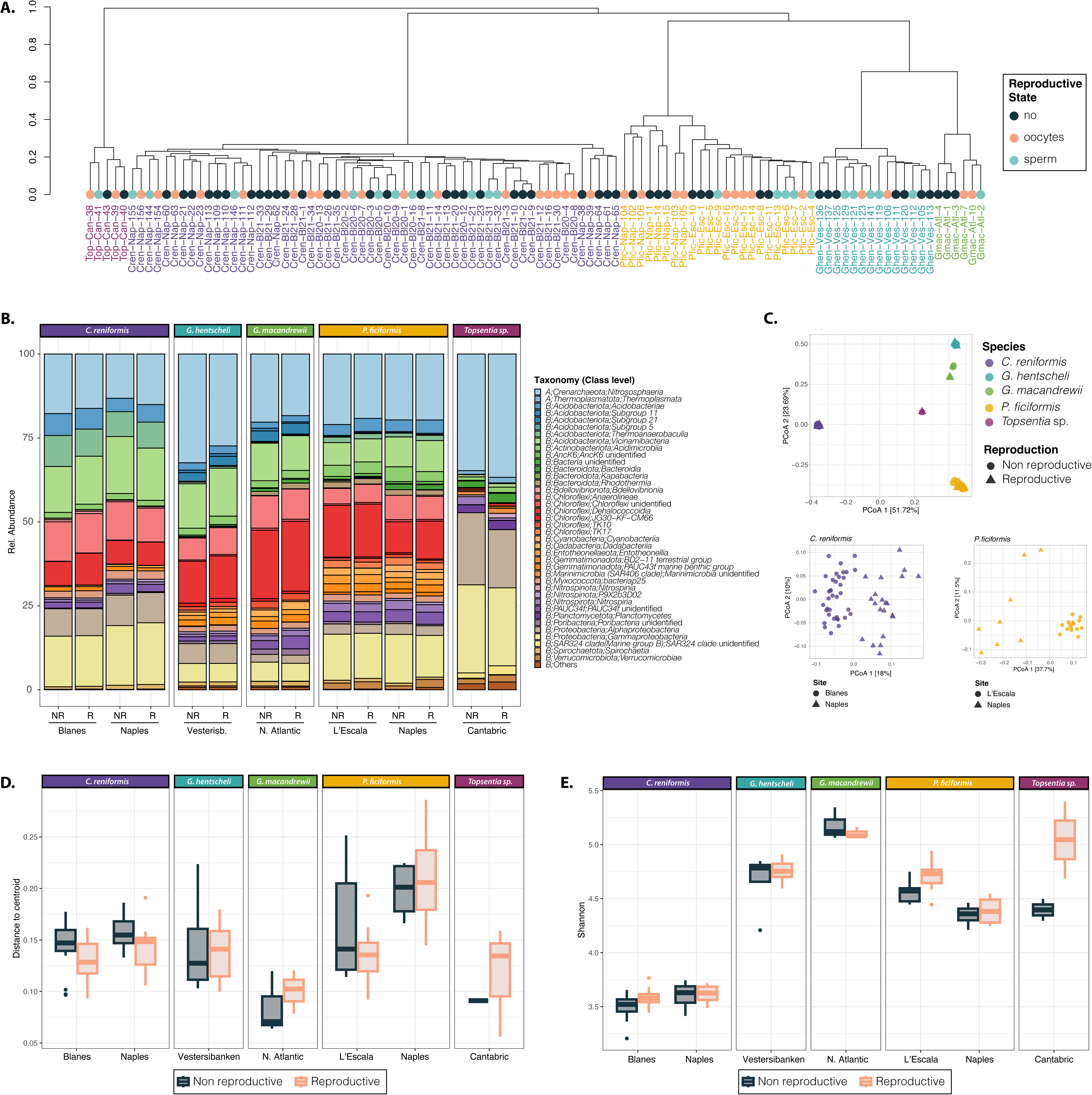
General microbiome results. **A**. Cluster dendrogram based on Bray-Curtis dissimilarity matrices of microbial communities of each sponge species. Each colour depicts a different sponge species their reproductive stage is indicated by different dot colors. **B.** Barplots showing the taxonomic composition at class level for each species, grouped by location and reproductive state. In the legend; A: Archaea, B: Bacteria. Taxa with relative abundances < 0.01 are grouped as “Others” **C.** Principal coordinate analysis (PCoA) plot based on Bray-Curtis dissimilarity index of the microbial composition across the different sponge species (indicated by different colours). Below, PCoAs of the two species with individuals in different locations (indicated by different shapes). Variation explained by the first two axes is indicated as %. **D.** Boxplot showing the Beta dispersion (distance to centroid) for each sponge species and location separated by reproductive state (reproductive or non-reproductive). **E.** Boxplot showing the Shannon diversity for each sponge species and location separated by reproductive state (reproductive or non-reproductive)

### Effect of reproductive state on sponge microbial diversity

Neither alpha diversity, assessed using Shannon index and richness, nor dispersion of the samples, measured as distance to centroid, showed statistically significant differences between reproductive and non-reproductive individuals for any species (Figure 3D, 3E, Supplementary stats). However, reproductive state impacted the beta diversity based on Bray-Curtis dissimilarity, for *G. macandrewii* and *P. ficiformis* from Naples (Supplementary stats, Figure 4C and 5C), but not for the other samples (Supplementary stats, Figure S1). In *G. macandrewii*, the ordination of samples revealed a distinct separation of individuals based on their reproductive state along the first axis, which accounted for 68.4% of the variation in microbiome composition (Figure 4C). Similarly, in the case of *P. ficiformis* from Naples, the ordination plot exhibited a clear segregation between reproductive and female individuals along the first PCoA axis, explaining 37.5% of the variation observed in the microbiome composition (Figure 5C).

**Figure 4.**
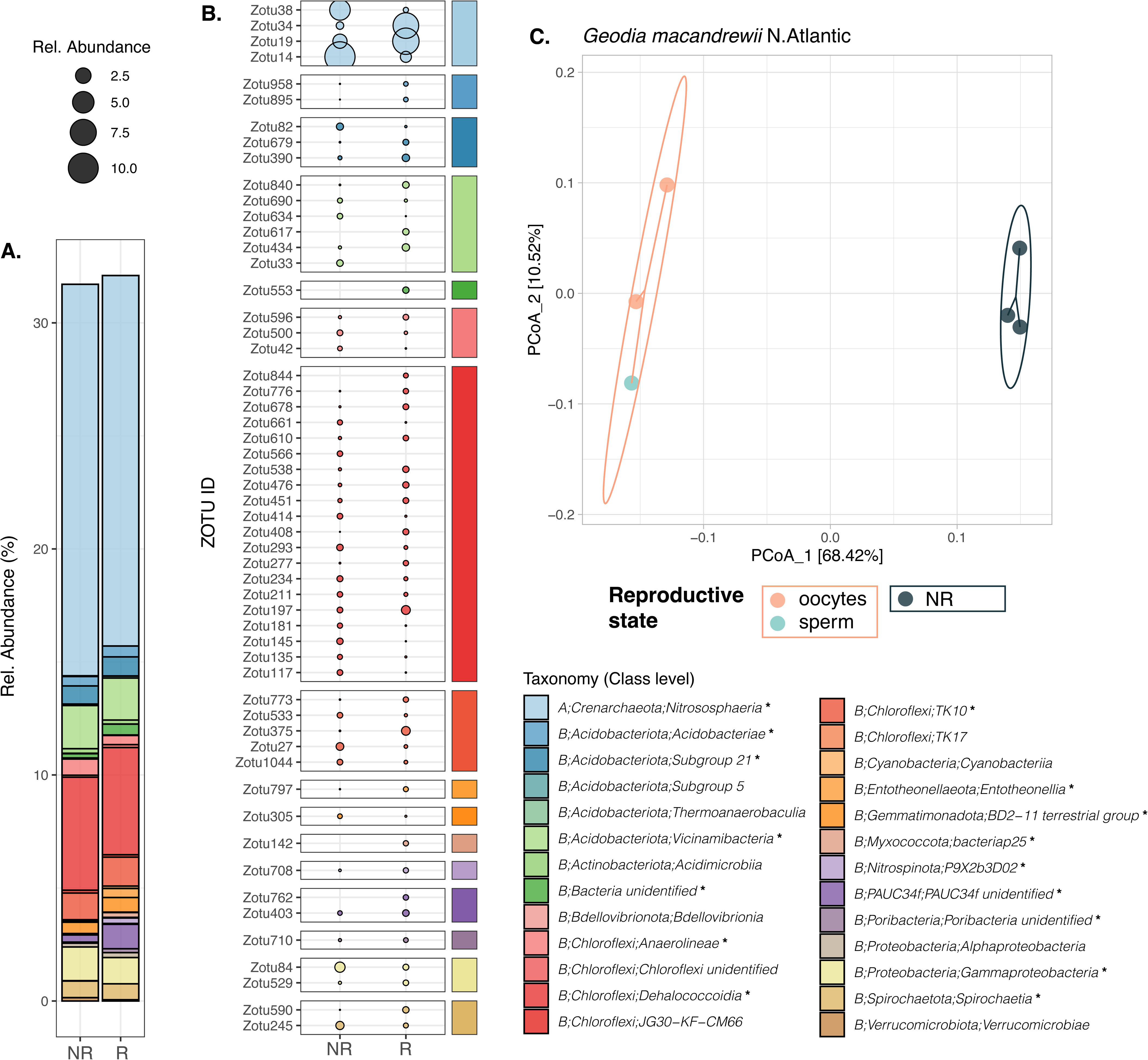
A. *Geodia macandrewii:* Barplot showing the taxonomic composition at class level of all the Differentially Abundant (DA) (see Table 2) ZOTUs in *G. macandrewii.* **B**. Average relative abundance of the most abundant DA ZOTUs (min. 0.2 avgRA in 1 sample) in reproductive and non-reproductive individuals. Taxonomic affiliation at class level of each ZOTU is indicated by colour bars. In the taxonomic legend, classes represented by * are the ones depicted in B. **C.** Principal coordinate analysis (PCoA) plot based on Bray-Curtis dissimilarity index of the microbial composition for *G. macandrewii*.

**Figure 5.**
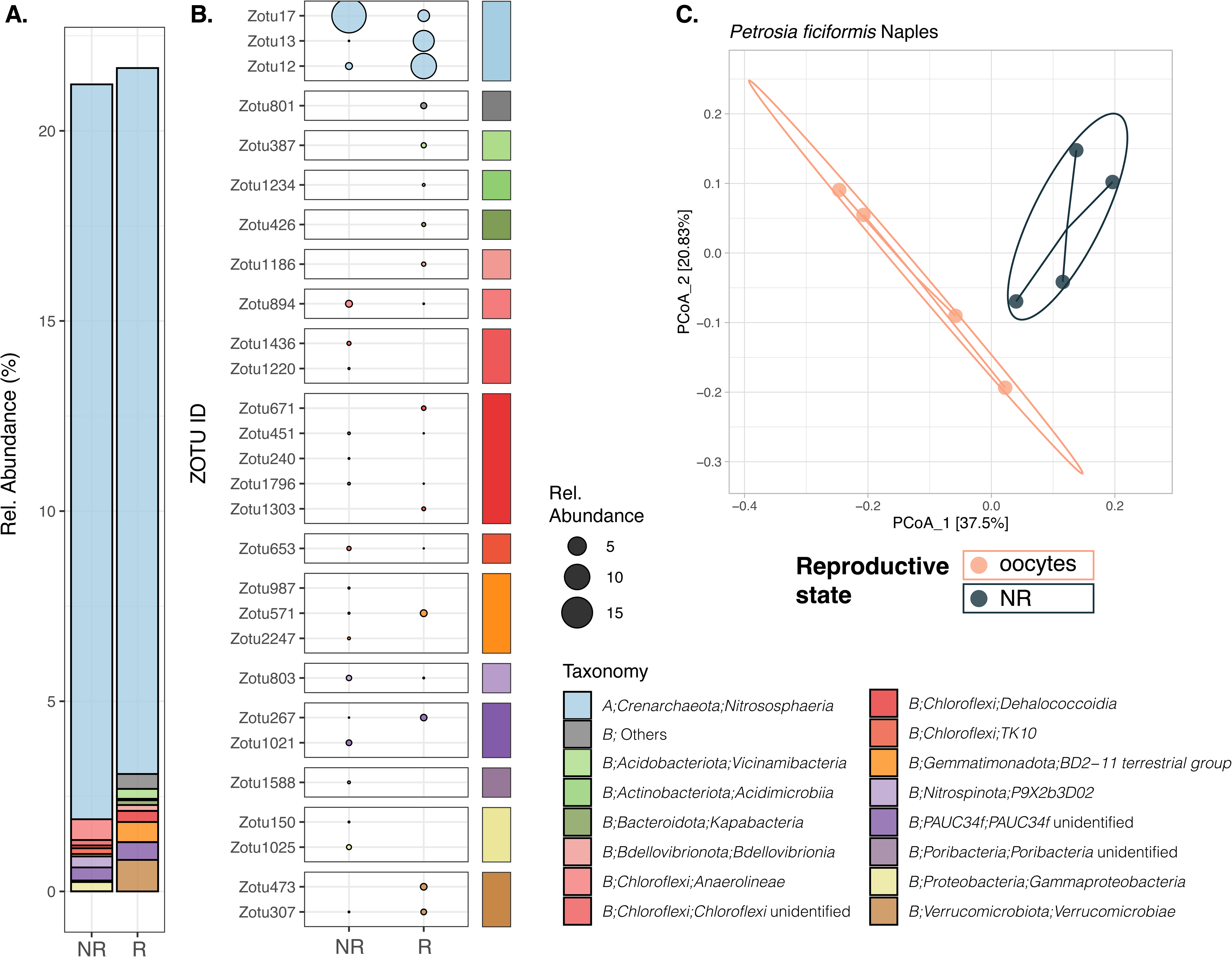
A. Barplot showing the taxonomic composition at class level of all the Differentially Abundant (DA) (see Table 2) ZOTUs in *P.ficiformis* from Naples. **B**. Average relative abundance of all the DA ZOTUs in reproductive and non-reproductive individuals. Taxonomic affiliation at class level of each ZOTU is indicated by colour bars. **C.** Principal coordinate analysis (PCoA) plot based on Bray-Curtis dissimilarity index of the microbial composition for *P.ficiformis* from Naples.

### Differential Abundance Analysis

Differential abundance (DA) analysis comparing reproductive and non-reproductive individuals, as well as females and non-reproductive individuals, yielded varying numbers of DA ZOTUs depending on the species (Table 2). In species such as *P. ficiformis* – Escala, *C. reniformis* from Blanes, *G. hentscheli* and *Topsentia* sp., significant enrichment of specific ZOTUs was detected in non-reproductive individuals. However, the number of these ZOTUs ranged from 2 to 24, and collectively, they represented minor fractions of the microbiome’s relative abundances accounting for less than 1% on average relative abundance (avgRA). In the case of *C. reniformis*-Naples 5 ZOTUs were found to be enriched in reproductive individuals, although they also represented small fractions of microbiome relative abundance (< 1% avgRA). In contrast, for *G. macandrewii* and *P. ficiformis –* Naples, a substantial number of 395 ZOTUs, representing 32% of avgRA of the sponge microbiome, and 28 ZOTUs, accounting for 22% of avgRA, respectively, displayed differential abundance between reproductive and non-reproductive individuals. A more in detailed explanation of these differences is presented below.

### Differentially Abundant ZOTUs in Geodia macandrewii

In the case of *G. macandrewii,* a striking pattern was observed: the three non-reproductive individuals exhibited significant enrichment of 239 ZOTUs compared to the reproductive individuals. Conversely, the reproductive individuals, comprising two females with previtellogenic oocytes (Figure 1A) and one male with sperm at the spermatogonial stage, displayed significant enrichment of 156 ZOTUs relative to the non-reproductive individuals (Table 2, Figure 4, Figure S2A). In the ternary plot (Figure S2 B), we observed numerous ZOTUs along with their relative abundances, represented by the size of the circles, positioned along the axes that correspond to reproductive individuals, with specific ZOTUs being mainly found in males or females. Similarly, the DA ZOTUs enriched in non-reproductive individuals were prominently depicted at the vertex associated with the non-reproductive stage. The DA ZOTUs were associated with 23 different phyla being *Crenarchaeota*, *Acidobacteria* and *Chloroflexi* the most abundant ones (Figure 4A). We calculated the proportion of ZOTUs identified as DA for each phylum and their contribution to the mean relative abundance of each phylum (Table S4). Notably, *Crenarchaeota* exhibited the highest contribution to the observed differences, accounting for a total of 16.8% average relative abundance (avgRA) of differentially abundant ZOTUs, representing 87% of the avgRA of the phylum. However, only 5 out of 38 *Crenarchaeota* ZOTUs were identified as DA. All of these ZOTUs belonged to the family *Nitrosopumilaceae*, with two of them (ZOTU 10 and ZOTU 19) classified as genus *Candidatus* Nitrosopumilus. Interestingly, ZOTU 10, ZOTU 19, and ZOTU 34 exhibited higher abundance in reproductive samples compared to non-reproductive individuals, while ZOTU 14 and ZOTU 38, were more abundant in non-reproductive individuals. Following *Crenarchaeota*, the phylum *Chloroflexi* contributed significantly to DA ZOTUs in terms of relative abundances, with 135 ZOTUs representing up to 7% of avgRA, encompassing 20% of the avgRA of the phylum. Within this phylum, the most dominant DA classes were *Dehalococcoidia*, *Anaerolineae*, and *TK10* cluster, and among them, some ZOTUs were more abundant in non-reproductive samples, while others showed higher abundance in reproductive individuals. A similar pattern was commonly observed within the other phyla, although relative abundances were generally lower. It is important to note that we did not observe any entire genus consistently being more abundant in one group than the other.

### Differentially Abundant ZOTUs in Petrosia ficiformis (Naples)

In the case of the sponge *P. ficiformis,* only the individuals collected in Naples exhibited a significant proportion of differential abundant (DA) ZOTUs between reproductive and non-reproductive individuals, in contrast to the individuals collected in L’Escala (Table 2). It is important to highlight that Naples’ samples were collected in July when the oocytes were still in early or mid-maturation stages, while the samples from L’Escala were collected in December, coinciding with the species’ reproductive peak when all oocytes were fully mature (Figure 1A). These differences in the reproductive stage may account for the disparate results observed within this species. However, it is worth considering that other factors such as the time of the year or specific location could also contribute to the variations between the two subsets for *P. ficiformis*. In total, 28 ZOTUs displayed significant differential abundances depending on the reproductive stage, with 15 ZOTUs enriched in non-reproductive individuals and 13 ZOTUs found at higher abundances in reproductive female individuals. These DA ZOTUs spanned across up to 13 different phyla. Much like the pattern observed in *G. macandrewii*, *Crenarchaeota* dominated the observed differences in *P. ficiformis*, contributing to a total of 19% average relative abundance (avgRA) across three differentially abundant ZOTUs. Remarkably, these ZOTUs accounted for 96% of the avgRA of the *Crenarchaeota* phylum within this species (Table S5). All the 3 DA *Crenarchaeota* ZOTUs were affiliated with the family *Nitrosopumilaceae* with ZOTU 12 and ZOTU 17 classified under the genus *Candidatus* Nitrosopumilus. Among these, ZOTU 12 and ZOTU13 displayed enrichment in reproductive individuals, while ZOTU17 exhibited higher abundance in non-reproductive individuals (Figure 5B). Importantly, it is noteworthy that the remaining phyla made up less than 1% of avgRA. Similar to our observations in *G. macandrewii*, there was no consistent pattern of an entire genus being consistently more abundant in one group over the other in *P. ficiformis*.

### Phylogeny of the most abundant Differentially Abundant ZOTUs

For both *G. macandrewii* and *P. ficiformis*, the most abundant taxa contributing to the disparities belonged to a single archaeal phyla *Crenarchaeota*, and the class *Nitrososphaeria* (Figure 4-5). The phylogeny of these ZOTUs found in those sponge species revealed that 4 ZOTUs with differential abundance between reproductive and non-reproductive samples fell into several clades with uncultured archaea, which hampered further taxonomic assignation (Figure 6). For the rest of differentially abundant ZOTUs found, one of them clustered with the newly described *Candidatus* Nitrosokoinonia, isolated from sponges, and the rest to *Nitrosopumilus*. Interestingly, in *G. macandrewii,* two DA ZOTUs in reproductive samples belonged to the clade containing both *Nitrosopumilus* and *Candidatus* Nitrosopumilus, clustering with other non-differential ZOTUs also present within *G. macandrewii*, and the other differentially abundant ZOTU belonged to a clade of uncultured taxa (Figure 6). Both differentially abundant ZOTUs found in the non-reproductive condition of *G. macandrewii* belonged to the same clade within the uncultured taxa (Figure 6). Similarly, the differentially abundant reproductive ZOTUs in *P. ficiformis* belonged to clade of uncultured taxa and to the clade of *Nitrosopumilus*, but the only differentially abundant ZOTU in the non-reproductive condition belonged to the newly erected genus *Candidatus* Nitrosokoinoia (Figure 6).

**Figure 6.**
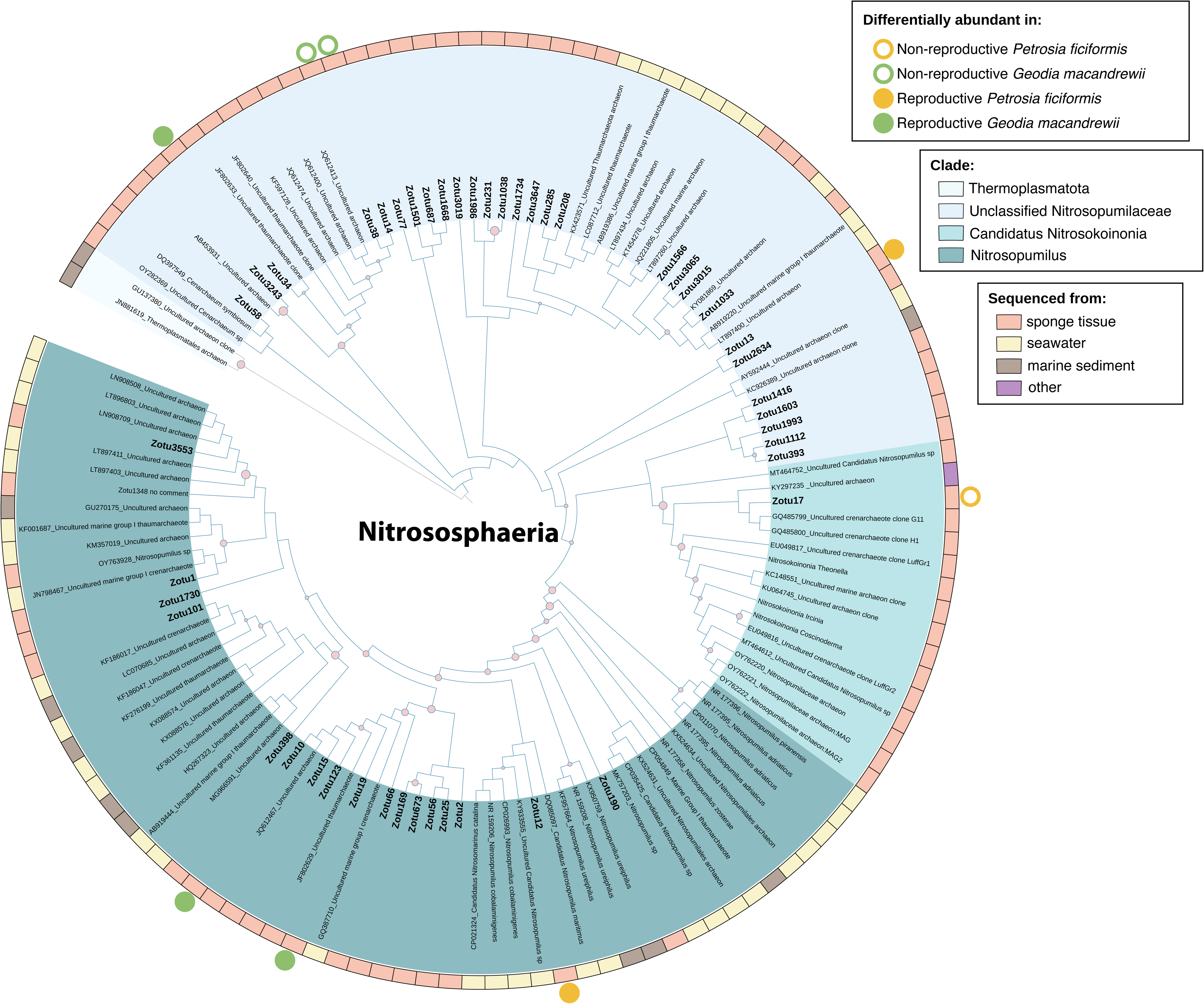
Phylogenetic tree for the Thermoplasmatota and Nitrososphaerota ZOTUs of *Geodia macandrewii* and *Petrosia ficiformis*. Bootstrap expectation values are shown as red circles (only over 0.7).

### Differential gene expression in Geodia macandrewii

We found a total of 345 differentially expressed transcripts between the reproductive (134 transcripts) and non-reproductive (211 transcripts) individuals of *G. macandrewii* (Figure 7A). Among those transcripts, only 24 and 28 were annotated in reproductive and non-reproductive individuals, respectively (Table S6). The reproductive condition (regardless of the sex of the sponges) showed overexpression of genes that included several associated to mitotic proliferation processes such as *G2 mitotic-specific cyclin-B1* (CCNB1), *Tyrosine protein kinase yes* (YES), *M-phase inducer phosphatase* (MPIP) and *Histone H2AX* (Table S6). The gene ontology (GO) enrichment performed with the upregulated genes in the reproductive condition revealed that an important proportion of them were associated to immune response and immune system processes, and included *60 kDa heat shock protein* (HSPD1), *Deleted in Malignant Brain Tumors 1 Protein* (DMBT1), *ADAMTS 2* (ADAMTS2 or ATL2), *TNF receptor-associated factor 5* (TRAF5), and *Indoleamine 2,3-dioxygenase 2* (I2302) (Figure 7B and FigureS3). The non-reproductive condition, in turn, showed upregulation of genes involved in ribogenesis and several metabolic pathways, such as the iron-sulfur cluster assembly, and lipid and amino acid metabolic pathways (Figure 7B).

**Figure 7.**
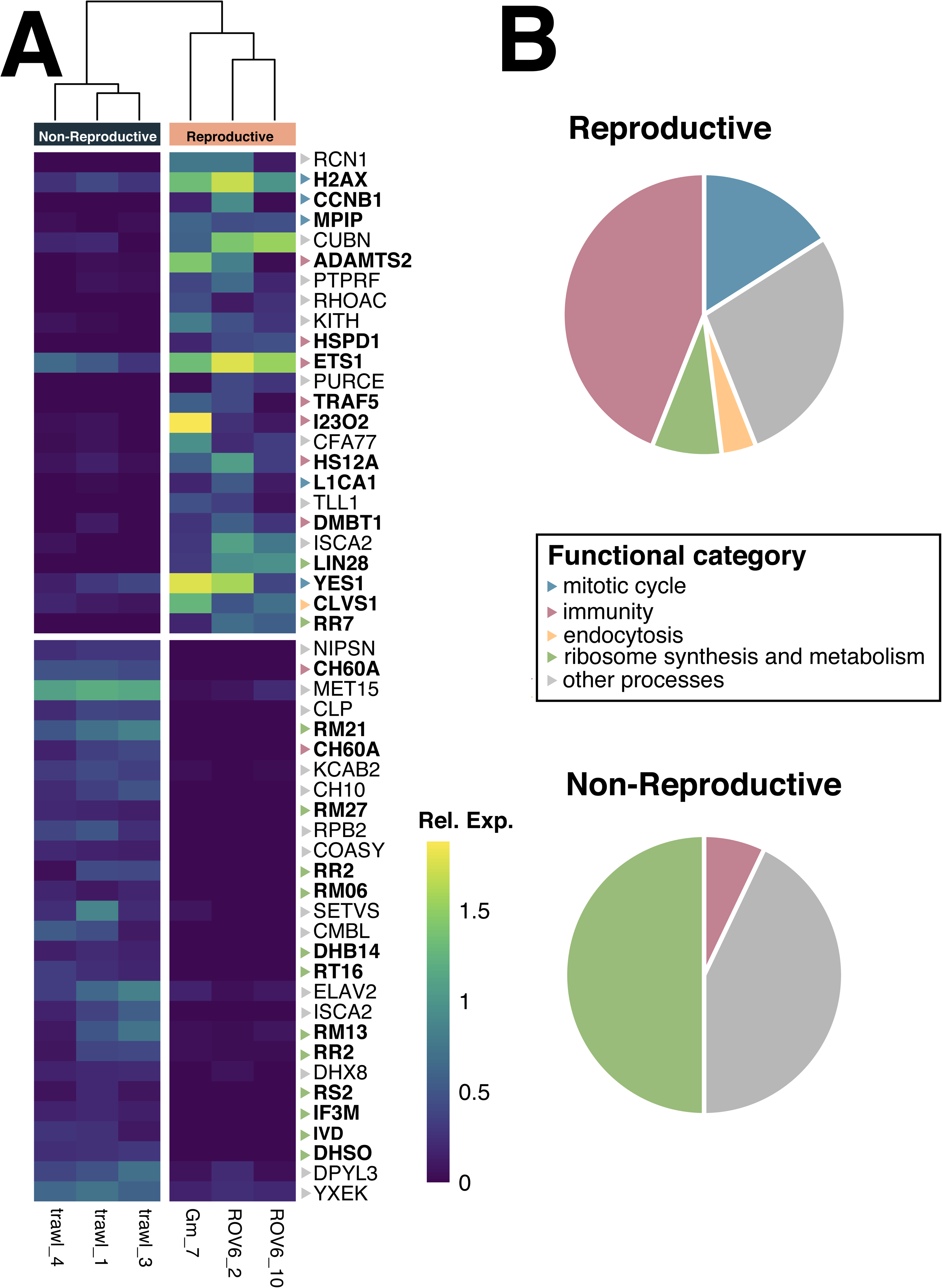
Transcriptional changes in *Geodia macandrewii*. **A**. Heatmap showing the expression values (log transformed) of the differentially expressed genes between reproductive and non-reproductive individuals. Only annotated genes are shown. The color scale for the genes represents their functional category. **B.** Pie chart depicting the major functional categories recovered in the reproductive and non-reproductive conditions.

## DISCUSSION

### Dynamics of the sponge microbiome during reproduction

During the reproductive period, most of the studied sponges exhibited high microbiome stability, characterized by highly species-specific microbial communities, a trait commonly observed in many sponge species (Thomas et al. 2016). The predominance of the class *Nitrosphaeria* among the studied species was anticipated, given that ammonia-oxidizing archaea (AOA) are crucial for the nitrification process in marine sponges and are widely distributed across various marine sponge species (Glasl et al. 2024). Our findings indicate that significant microbiome differences between reproductive and non-reproductive individuals were observed only in sponges with oocytes in early phases of maturation (previtellogenic oocytes), when the vitellogenesis is taking place. This suggests that these initial stages may be the critical point where microbial changes are more likely to occur because of their role in the process. Given that most of the species were collected past this point, a temporal study tracking the same individual throughout their reproductive cycle would be necessary to collect samples at the same state in the different species in order to assess this hypothesis on the role that the microbiome plays during gametogenesis.

In the case of the sponges with previtellogenic oocytes, i.e. *Geodia macandrewii* and *Petrosia ficiformis*-Naples, such changes were primarily driven by differential abundances of specific ZOTUs between reproductive and non-reproductive individuals. In both species, ZOTUs belonging to the class *Nitrosphaeria* were by far, the most abundant ones and contributed significantly to these disparities. Interestingly, among them, some differential taxa belonged to *Nitrosopumilus* and a clade with uncultured archaea, while in other species the differential ZOTU was affiliated to the newly described nitrifying symbiont lineage *Candidatus* Nitrosokoinonia. Members of this new genus rely on endogenously produced nitrogenous compounds by the host and have been found across many sponge taxa, but noteworthy is the abundance in the larvae of the sponge *Coscinoderma matthewsi*, which suggests that these archaeal symbionts are vertically transmitted and might play crucial roles in the early life stages of sponges (Glasl et al. 2024). The importance of archaea for reproduction was highlighted previously in a paper by Engelberts et al. (2022), where they reported that the crenarchaeote *Candidatus* Nitrosospongia ianthellae was the dominant taxon in the oocytes and embryos although not in the adult sponges, and it exhibited much less relative abundance in male specimens than in the rest of samples. In addition, they confirmed that *Candidatus* Nitrosospongia ianthellae was vertically transmitted from the sponge mother to the offspring (Engelberts et al. 2022). Even though the relative abundance of such taxon in non-reproductive adults was not evaluated, it underscores the importance of archaeal symbionts for sponge reproduction, with variability across sexes and life stages that could potentially impact the functional roles provided to the sponge host.

Our results they show a divergent trend of abundance in archaeal taxa during reproduction that is worth discussing, although they are based on qualitative data (relative abundance), and absolute quantitative data would be necessary to confirm the increase or decrease of specific microbes. We propose two possible roles of these archaeal taxa. First, certain microbes and not others may support the high energetic demands during the initial phases of oogenesis and the production of substantial yolk quantities during the vitellogenic phase of oogenesis. This would suggest a mechanism where the sponge selectively digests specific microbial symbionts, utilizing a finely tuned recognition system to target the most nutritionally valuable microbes from their microbiome. Evidences of symbiont phagocytosis has been widely reported (e.g. Saleuddin et al. 2024; Maldonado and Riesgo 2009; Lanna and Klautau 2012; Leys et al. 2018; Koutsouveli et al. 2018), indicating that symbiont digestion is balancing the carbon and nitrogen budget, at least for certain species (Leys et al. 2018). Here, we documented symbiont phagocytosis in all of our studied sponges, especially during their reproductive periods. Whether this phagocytosis is carried out selectively, however, remains to be elucidated. In this sense, sponge-associated *Nitrososphaerota* (previously known as *Thaumarchaeaota*) may use eukaryotic-like proteins and modified surface layer glycoproteins in their membranes, which are absent from free-living relatives, to avoid digestion from the sponge host (Moeller et al. 2019; S. Zhang et al. 2019; Haber et al. 2021). Since it is unlikely that the mechanism for symbiont digestion facilitation appears in the reproductive individuals as a result of changes in symbiont physiology, the physiological state of the sponge (completely altered compared to individuals that are not reproducing) seems a more likely mechanism behind the alteration in symbiont digestion (see sections below). The second possible role of these microbes would be triggering oogenesis, where a significant change in their abundance—whether an increase or decrease—could promote the onset of oogenesis. Both of these scenarios have been previously suggested for other animal species. For instance, in *Hydra viridis,* its *Chlorella* symbionts are believed to play an oogenesis promoting role (Habetha et al. 2003). The authors suggest that while the nutrients provided by the photosynthetic algae may support the energetic demands of oogenesis, the inability to produce eggs in the absence of symbionts points to a triggering role of these symbionts. Another example of microbial dependence for reproduction is seen in the moon jellyfish *Aurelia aurita* (Weiland-Bräuer et al. 2020), where the generation of daughter polyps depends on the presence of the native microbiota. Although molecular evidence is lacking, key bacterial OTUs correlated with the release of ephyrae, suggesting fundamental roles for certain bacterial metabolites in this process (Weiland-Bräuer et al. 2020). Additionally, specific bacteria are known to interfere with the sexual reproduction of a benthic diatom, where the presence of *Maribacter* sp. reduces reproductive success by decreasing the production of a sexual attraction pheromone, and presence of *Rosevarius* sp. enhances reproduction by increasing the expression of enzymes that synthesize pheromone precursors (Cirri et al. 2019). Moreover, in chickens, certain reproductive tract microbial species are strongly associated to enhanced egg production and are related to immune functions (Su et al. 2021). In the case of *Drosophila melanogaster,* the microbiome has been shown to influence life history strategy by affecting the balance between early reproduction and somatic maintenance (Walters et al. 2020). The authors hypothesize that certain bacteria may produce key metabolites that play a crucial role in female fecundity. All of the above-mentioned examples support the idea that reproductive microbiomes can significantly influence reproductive function and performance, as reviewed by Rowe et al. (2020), and sponge microbiomes might not be an exception to this respect.

### Reproductive Microbiomes in sponges: exclusive to a few species?

The majority of the specimens analysed in this study belong are HMA sponges, known for harbouring dense and stable microbial communities within their tissues (Hentschel et al. 2003; Moitinho-Silva et al. 2017). Notably, the ability of HMA sponges to recruit high symbiont abundances has been associated with evolutionary shifts towards gonochoristic strategies (Riesgo, Novo, et al. 2014; Díez-Vives et al. 2022). Therefore, it has been suggested that microbes might play a crucial role in supporting the reproductive processes of these sponges (Díez-Vives et al. 2022). In males, they likely fulfil nutritional demands essential for spermatogenesis to progress, particularly when filtration is compromised during the transformation of choanocyte chambers into spermatic cysts (Riesgo and Maldonado 2008). Conversely, in females, microbial communities may contribute to the high nutritional requirements for yolk formation during oogenesis. Notably, all HMA species analysed in this study exhibit gonochorism and oviparity, albeit with variations in their gametogenic periods, which might differently affect their nutritional needs (Riesgo and Maldonado 2008). For instance, *C. reniformis* displays a short oogenic period of three months and spermatogenesis lasting one to two weeks. Reports indicate that its spermatic cysts, though small, occupy a significant portion of the sponge mesohyl, potentially limiting its feeding capacity (Riesgo and Maldonado 2008). In contrast, *P. ficiformis* has a longer oogenic period lasting about seven months, with spermatogenesis taking approximately 2.5 weeks (Maldonado and Riesgo 2009). As for the *Geodia* species, their reproductive period spans more than three months, with oocytes containing a high lipid yolk content acquired through autosynthesis during three months and sperm developing for one to three months in cysts (Koutsouveli et al. 2020). Although both *Geodia* species invested a similar proportion of resources in spermatic cyst production (∼3%), there was a significant disparity in the number of oocytes present, with *G. macandrewii* housing approximately 116 million oocytes per individual compared to the 4 million reported in *G. hentscheli* (Koutsouveli et al. 2020). This stark difference in fecundity between the *Geodia* spp. could point to a much larger energetic investment in *G. macandrewii* for which the microbiome might be even more crucial during the reproductive period than for *G. hentscheli*.

In addition to the duration of gametogenesis and the proportion of mesohyl occupied by male or female gametes, the stage of gametogenic development likely influences the nutritional demands of the sponge. Extensive research has been dedicated to understand how ingested food resources are partitioned among competing metabolic processes, especially between growth and reproduction (Powell, Marsh, and Watts 2020). In particular, several processes of the gametogenic cycle are more energetically demanding than others (Powell, Marsh, and Watts 2020). Among them, the vitellogenesis stands out because of the rapid carbohydrate metabolism, protein turnover and lipid production occurring to allocate reserves to oocytes as yolk platelets (Eckelbarger and Hodgson 2021). Usually in invertebrates, ovaries and developing oocytes exhibit lipid profiles that directly reflect their dietary sources (e.g., (Marsh et al. 1990)). Among the species examined in this study, most exhibited vitellogenic oocytes nearing completion of maturation, suggesting that their nutritional requirements were likely adequately met. However, specimens of *G. macandrewii* and *P. ficiformis* (from Naples), contained previtellogenic oocytes in their mesohyl, indicating that they were collected during periods of high nutritional demand, with maturation percentages below 50%. These sponges presented differences in their microbiomes during reproduction compared to non-reproductive specimens, mostly affecting class *Nitrosphaeria* members. Why these two species were selectively consuming large quantities of specific symbionts is still a matter for further research, but the reasons might likely be related to the energetic opportunities that these symbionts embody more than others within the mesohyl of the sponges. Our results underscore the importance of considering the specific stage of gametogenesis when assessing the nutritional needs and the potentially associated microbiome changes for different sponges’ species in reproductive contexts, since most remarkable changes might likely be associated to transient, but energetically demanding short periods of time.

### Sex-specific differences at the microbiome level

To our knowledge, microbiome sex-specific differences in sponges have not been evaluated properly, primarily due to the challenges associated with identifying males and females within a population, which involves preparing histological sections of the samples. In addition, oogenesis typically takes relatively long periods, whereas spermatogenesis is usually very rapid, spanning only a few weeks, and therefore, detecting males during a single sampling event is less likely. The only exception though, is the study by Engelberts et al. (2022), who found significant differences in the microbial composition of females and males of *Ianthella basta*, although the differences were not further evaluated. Despite we did not have enough replication to properly detect significant differences between males and females in our study species, in an exploratory analysis, specific differentially abundant ZOTUs were detected for both males and females in *G. macandrewii* (Figure S2B). Our results thus suggest the possibility that certain microbiome changes do occur during spermatogenesis, but whether these occur as a result of specific physiological demands of sperm formation, such as glycogen accumulation, or associated to sex determination processes is impossible to test here.

Sex-specific differences at the microbiome level have been reported in many other animals (Bates et al. 2023). For instance, in the octocoral *Lobophytum pauciflorum*, although minor differences between sexes were detected at the microbial class level, significant differences at the OTU level were observed. Females exhibited a two-fold higher relative abundance of unassigned sequences compared to males, suggesting that novel sequences may contribute to the observed sex differences (Wessels et al. 2017). The mucous layer in the cephalopod *Octopus vulgaris* exhibits a sex-specific symbiosis in which microbes benefit from easy access to distinct substrates present in female and male skin, respectively (Rodríguez-Barreto et al. 2024). Another example is the nematode *Caenorhabditis elegans*, where bacteria exhibit sex-specific effects, leading to differential developmental rates in males and females (Santos et al. 2023). While sex-specific microbiomes could be the result of sexual size dimorphism or to sex-specific differences in habitat selection (Bates et al. 2023), different physiological demands of the reproductive process in each sex might also be behind the divergence in their microbiomes. Microbial sex differences may have implications for population adaptation to local microbial communities in nature, which should be considered in any study involving gonochoristic (male and female) species.

### Sponge immunity modulates the reproductive microbiome

The sponge innate immune repertoire is complex and includes homologues of many pattern recognition molecules and components of signal transduction pathways (Riesgo, Farrar, et al. 2014; Degnan 2015; Pita et al. 2018), whose functions in vertebrates have been extensively studied (Degnan 2015). The immune system then plays a critical role in mediating the animal–bacterial crosstalk needed for finely tuned discrimination between microbes of various relationships in a context of high densities of symbiotic microbiomes (Degnan 2015; Pita et al. 2018; Schmittmann, Franzenburg, and Pita 2021). While there is extensive literature regarding the transmission of the microbiota from the maternal sponge to the offspring (reviewed in Carrier et al. 2022; Díez-Vives et al. 2022), very little is known about the dynamics of the microbiome during the gametogenic processes in sponges and how that is modulated. The intimate relationship between the immune and the reproductive systems is observed across the eukaryotic tree of life, and for most animals, the use of the immune system is costly and thus must be traded off against other traits, such as survival and reproduction, to balance an organism’s investment into all the traits needed to promote evolutionary fitness (Barribeau and Otti 2020). In the case of complex holobionts though, the immune system helps to modulate the microbiota to favor reproduction (Al-Nasiry et al. 2020). Here, we found the upregulation of several genes of the innate immune system of *G. macandrewii* during its gametogenic cycle, pointing to an important role of the immune response in the modulation of the microbiome during reproduction, which we hypothesize to be involved in targeting and phagocytosing symbiotic microbes to enhance nutrition. The sponge host innate immune system includes a complex repertoire of immune receptors, so-called pattern recognition receptors (PRRs), that recognize the microbe associated molecular patterns (MAMPs) present in prokaryotes but absent in eukaryotes (Janeway and Medzhitov 2002). MAMPs include components of bacterial cell walls and membranes, lipopolysaccharides (LPS), but also endogenous ligands derived from damaged cells (Yu, Wang, and Chen 2010). PRR stimulation ultimately leads to the production of inflammatory cytokines or antimicrobial peptides (AMPs) (Brennan and Gilmore 2018), or the activation of cellular immune effectors such as phagocytes. Here we found over expression of the PRR glycoprotein Deleted in Malignant Brain Tumors 1 (DMBT1), which is known as agglutinin or salivary protein in mammals, and is a member of the scavenger receptor (SRCR) superfamily, present in all metazoans (Ligtenberg, Karlsson, and Veerman 2010). DMBT1 binds to several Gram+ and Gram− bacteria based on a variable motif, producing agglutination (Bikker et al. 2004; Madsen, Mollenhauer, and Holmskov 2010). *Geodia macandrewii* had over 72 genes encoding for DMBT1, but only one was upregulated during the reproduction (Figure 7), and could be potentially involved in targeting specific symbionts to be phagocytosed. While in humans most microbial phagocytotic processes are carried out by macrophages, sponges have archaeocytes and choanocytes, which are the most abundant cells in the sponge body. Macrophage activation is tightly regulated, and both TRAF and ADAM proteins contribute to this process (Hofer et al. 2008; Zhu, Yue, and Wang 2014; Lalani et al. 2018). The upregulation of TRAF5 and ADAMTS2 genes in *G. macandrewii* suggests that phagocytotic process are activated during reproduction (Figure 7). In other sponges, TRAF5 changes induced by environmental challenges indicate a strong role regulating antimicrobial, inflammatory, and apoptotic mechanisms, by modulating their ability for symbiont recognition and pathogen clearance (Posadas et al. 2022).

Several other genes with various functions in the animal immune system were upregulated during the reproduction of *G. macandrewii* such as I23O2 (also known as IDO in humans) and HSPD1. The products of these genes are known for their roles in immunoregulation not only coordinating inflammation and innate immunity, but also additionally establishing self-tolerance in humans (Wu et al. 2018). For instance, during human reproduction, uterine macrophages maintain a tolerogenic and receptive microenvironment, by producing interleukin-10 (through activation of HSPD1), TGF-β, and I23O2 (Swartwout and Luo 2018; Al-Nasiry et al. 2020). IDO (or I23O2) contributes to ‘metabolic immune regulation’ by catalyzing oxidative catabolism of the essential amino acid tryptophan (TRP) along the kynurenine (KYN) pathway and is primarily expressed by macrophages (Munn and Mellor 2013). TRP degradation was described as an innate immune mechanism of host defense against infection, and therefore, it is understood that IDO contributes to immune regulation via local metabolic changes in the immediate microenvironment and local tissue milieu (Munn and Mellor 2013). Recently, Indoleamine 2,3-dioxygenase activity has been directly related to regulation of gut microbiota, specifically determining its composition (Laurans et al. 2018). Finally, HSPD1 encodes for the chaperokine Hsp60, recognized by receptors of both the innate and adaptive immune systems, and when the levels of Hsp60 are high pro-inflammatory responses are initiated, and when they decrease, there is an activation of anti-inflammatory responses (Zininga, Ramatsui, and Shonhai 2018; Quintana and Cohen 2011).

## CONCLUSIONS

Overall, it seems that a complex interplay between nutritional needs and the immune system of the sponge host influence the composition of the microbiome during reproduction, targeting and phagocytosing specific taxa to gain specific metabolites and aid in the yolk production and energy expenditure of the sponge. In the middle of these host physiological changes, archaeal class *Nitrosphaeria* emerges as a key player in the host nutrition during yolk production. Our results indicate that host sexual systems can greatly influence the microbiota, and exemplify why assessing sex and reproductive conditions of sponges can be of crucial importance to understand the evolutionary and ecological trajectories of sponge-microbe associations.

## Acknowledgments

We are indebted to Xavier Turon, Konstantina Mitsi, Laura Núñez-Pons, Patricia Álvarez-Campos, Javier Cristobo, Pilar Ríos, Hans Tore Rapp, Francisca Carvalho, Paco Cárdenas, and Manuel Maldonado, for sending specimens to study and help during field sampling. AR acknowledges funding from the Spanish Ministry of Science and Innovation, grants RYC2018-024247-I and PID2019-105769GB-I00, both funded by MCIN/AEI/10.13039/50110001103 and EI “FSE invierte en tu futuro”. CDV received the support of a fellowship from “la Caixa” Foundation (ID 100010434), with the fellowship code LCF/BQ/PI22/11910040. MT was supported by a JdC (Juan de la Cierva Formación, 2020) personal grant (FJC2020-043677-I). ST received funding from the grants PID2020-117115GA-100 of the Spanish Ministry of Science and Innovation and CNS2023-144572, and by the Ramón y Cajal grant RYC2021-03152-I, funded by the MCIN/AEI/10.13039/501100011033 and the European Union «NextGenerationEU»/PRTR». AV was supported by a European Union NextGenerationEU/PRTR grant (IJC2020-045256). Finally, part of this research was supported through the SponBIODIV project (AR and ST), a 2021-2022 BiodivProtect joint call for research proposals, under the Biodiversa+ Partnership co-funded by the European Commission, and with the funding organisations ‘Fundación Biodiversidad’ and FORMAS.

## Data availability

Raw sequence data are available from the NCBI SRA under project PRJNA1130701 (http://www.ncbi.nlm.nih.gov/bioproject/1130701) and metadata information is found in Table S1.

## Supplementary Figures

**Figure S1. Principal coordinate analysis (PCoA) plots** based on Bray-Curtis dissimilarity index of the microbial composition separated for each species and location, coloured according to their reproductive stage. Plots for *G. macandrewii* and *P.ficiformis* from Naples are shown in Figure 4 and 5 in the main text. Variation explained by the first two axes is indicated as %.

**Figure S2: *G. macandrewii.* ZOTUs A. Heatmap** of the most abundant differentially abundant (DA) ZOTUs between reproductive and non-reproductive individuals of *G. macandrewii,* with log transformed abundances represented in the colour temperature bar. Microbial ZOTUs are organized according to a hierarchical clustering based on Bray-Curtis Dissimilarity matrices. Sponge individuals (x-axis) are coloured according to Reproductive stage. Row names indicate the taxonomy of the DA ZOTUs (A: Archaea, B: Bacteria), in bold the 4 most abundant DA ZOTUs. **B. Ternary plot** of the ZOTUs distribution along reproductive stages (oocytes, sperm, non-reproductive) for *G. macandrewii.* Each circle represents a ZOTU and its size is proportional to its mean relative abundance across all samples. Only DA ZOTUs are coloured according to their taxonomy at class level.

**Figure S3. Treemap of the upregulated functions in reproductive individuals.** Size of the squares is proportional to the TMM expression values of each function.

## Supplementary Tables

**Table S1. Information table** on the samples used in the study with their information on species, collection site, date, and reproductive state.

**Table S2. Taxonomic composition at Phylum** level for each species ordered in descending mean total abundance (last column). Taxa with relative abundances < 0.01 are grouped as “Others”. Cr: *Chondrosia reniformis*, Gh: *Geodia hentscheli*, Gm: *Geodia macandrewii*, Pf: *Petrosia ficiformis*, Tp: *Topsentia* sp. In Taxonomy column, A: Archaea, B: Bacteria.

**Table S3. Taxonomic composition at Class level** for each species ordered in descending mean total abundance (last column). Taxa with relative abundances < 0.01 are grouped as “Others”. Cr: *Chondrosia reniformis*, Gh: *Geodia hentscheli*, Gm: *Geodia macandrewii*, Pf: *Petrosia ficiformis*, Tp: *Topsentia* sp. In Taxonomy column, A: Archaea, B: Bacteria.

**Table S4. Differentially abundant (DA) phyla detected for *G. macandrewii***. Columns 2 and 3 represent number of ZOTUs. While columns 5 and 6 represent average relative abundances. The proportion of DA ZOTUs (column 4) and their contribution to the avgRA of each phylum (column 7) are shown. In Taxonomy column, A: Archaea, B: Bacteria.

**Table S5. Differentially abundant (DA) phyla detected for *P. ficiformis* (Naples)**. Columns 2 and 3 represent number of ZOTUs. While columns 5 and 6 represent average relative abundances. The proportion of DA ZOTUs (column 4) and their contribution to the avgRA of each phylum (column 7) are shown. In Taxonomy column, A: Archaea, B: Bacteria.

**Table S6. Differential gene expression (DE) in *G. macandrewii.*** Table showing the DE transcripts between reproductive and non-reproductive individuals in *G. macandrewii* with their corresponding gene annotations and their expression levels in each sample.

